# Armoring STEAP1 CAR T cells with IL-18 potentiates antitumor activity in Ewing sarcoma

**DOI:** 10.64898/2025.12.08.693072

**Authors:** Vipul Bhatia, Annabelle Tsao, Truman Chong, Peter. P Challita, Keke Liang, Erolcan Sayar, Jiaoti Huang, Elizabeth R. Lawlor, Michael C. Haffner, Behnam Nabet, John K. Lee

## Abstract

**Background:** Ewing sarcoma (EwS) is a highly aggressive cancer driven by the EWS::FLI1 fusion oncoprotein affecting children, adolescents, and young adults. Six transmembrane epithelial antigen 1 (STEAP1) is a cell surface antigen transcriptionally controlled by EWS::FLI1 that is broadly expressed in EwS, positioning it as a rational immunotherapy target. However, translating CAR T therapy to solid tumors requires overcoming barriers to potency while maintaining safety.

**Methods:** Analyses of transcriptome and proteome data were performed to evaluate the effects of EWS::FLI1 perturbation on STEAP1 expression at the transcript and protein levels in EwS models. STEAP1 expression was validated in EwS patient tissues by immunohistochemistry. Second-generation STEAP1-BBζ CAR T cells were tested in orthotopic and disseminated EwS xenograft models. To enhance antitumor activity, an IL-18-armored STEAP1 CAR was engineered. Dose-dependent therapeutic efficacy and safety were evaluated through measurement of tumor burden, survival, and observation for gross toxicities.

**Results:** STEAP1 was expressed in ∼97% of primary EwS tumors and directly associated with EWS::FLI1 fusion protein expression in EwS cell lines. In orthotopic EwS models, STEAP1 CAR T cells induced complete tumor regression at 5 x 10^6^ cells. In disseminated disease models, responses were dose-dependent with no evidence of antigen loss. Notably, IL-18 armored STEAP1 CAR T cells achieved complete responses in ∼80% of mice at a reduced dose of 10^6^ cells without overt toxicity.

**Conclusions:** These data establish STEAP1 as a clinically relevant and highly expressed target in EwS and demonstrate that IL-18 armoring significantly improves CAR T cell efficacy by enhancing potency evident through antitumor activity at reduced cell dose. STEAP1 CAR T cells are currently under evaluation in a first-in-human phase 1/2 dose-escalation clinical trial for metastatic castration-resistant prostate cancer (NCT06236139) and these studies support future clinical translation of STEAP1 CAR T cell therapy for relapsed/refractory EwS.

## BACKGROUND

Ewing sarcoma (EwS) is an aggressive malignancy of bone and soft tissue that primarily affects children, adolescents, and young adults. Histologically, EwS is characterized by sheets of small, round, blue cells with strong membranous CD99 expression. Molecularly, ∼85% of EwS cases harbor the chromosomal translocation t(11;22)(q24;q12), which results in the fusion of EWSR1 with a member of the E26 transformation-specific (ETS) transcription factor family, most commonly FLI1, producing the oncogenic EWSR1:FLI1 gene fusion^1–3^. These pathognomonic fusion drives tumor initiation and sustains the malignant phenotype of classical EwS.

Despite risk-adapted multimodal therapy including intensive chemotherapy, surgery, and radiotherapy, outcomes for patients with metastatic or recurrent EwS remain poor with long-term survival rates below 30%^4,5^. This highlights the urgent need for more effective treatments. Immunotherapy, particularly chimeric antigen receptor (CAR) T-cell therapy, offers a promising avenue. By genetically redirecting T cells to recognize tumor antigens independently of major histocompatibility complex (MHC)-T cell receptor (TCR) signaling, CAR T cells have revolutionized treatment in hematologic malignancies^6,7^. However, translating this success to solid tumors like EwS remains challenging due to issues such as antigen heterogeneity, on-target off-tumor toxicity, immunosuppressive tumor microenvironments (TME), and limited T cell persistence and trafficking^8–10^. Nevertheless, early-phase clinical trials targeting disialoganglioside (GD2)^11–13^ and B7-H3 (CD276)^14–16^ have demonstrated initial efficacy and safety of CAR T cell therapies in solid tumors affecting pediatric and adolescent and young adult populations such as neuroblastoma and diffuse midline glioma supporting the feasibility and promise of this approach in solid malignancies.

One emerging immunotherapeutic target for solid tumors is six-transmembrane epithelial antigen of the prostate 1 (STEAP1), known to be overexpressed in multiple malignancies including prostate cancer, non-small cell lung cancer, and EwS^17,18^. STEAP1 is a metalloreductase that facilitates iron and copper transport and forms homo- or hetero-trimeric complexes with STEAP family members^19,20^. In EwS, STEAP1 is a direct transcriptional target of the oncogenic EWS::FLI1 fusion protein^21^ which drives STEAP1 expression through binding to GGAA-microsatellite elements. While EWS::FLI1 remains difficult to target pharmacologically due to its nuclear localization and role as a transcription factor, restricted STEAP1 surface expression in normal tissues^22^ and high STEAP1 expression in tumors make STEAP1 a more practical and clinically tractable therapeutic target. In multiple tumor types including, STEAP1 has been shown to promote proliferation, immune evasion, and maintenance of oncogenic transcriptional programs^21^.

In recent years, multiple immunotherapeutic strategies targeting STEAP1 have emerged. Among these strategies, T cell receptor (TCR)-engineered T cells specific for STEAP1-derived peptides presented on human leukocyte antigen HLA-A*02:01^23,24^ have shown encouraging preclinical activity against EwS models. Though they show effective preclinical activity, their clinical translation remains constrained by HLA restriction and tumor immune editing. T cell engagers (TCE) engineered to bind CD3 and STEAP1 have also demonstrated substantial antitumor activity in preclinical prostate cancer and EwS models^25,26^. The most clinically advanced TCE targeting STEAP1 under investigation is xaluritamig (AMG 509)^26^ which has shown highly encouraging activity in a phase I clinical trial in heavily pretreated metastatic castration-resistant prostate cancer (mCRPC) patients, achieving a 50% decline in serum prostate specific antigen (PSA50 response rate) of 59% and an objective response rate of 41% at higher doses with manageable cytokine release syndrome as the most common adverse event^27^. While challenges such as limited serum half-life and T cell exhaustion may impact the durability of TCE therapies, these early clinical results underscore the therapeutic promise of STEAP1-directed T cell redirection. Building on this, we previously developed a second-generation STEAP1-targeted CAR T cell therapy which demonstrated robust antitumor activity and a favorable safety profile in both immunodeficient and immunocompetent murine models of prostate cancer^28^. This autologous STEAP1 CAR T cell therapy is currently being evaluated in a first-in-human phase I/II clinical trial in mCRPC (NCT06236139).

In this study, we profiled STEAP1 expression across various pathological stages of EwS to better characterize its range of expression within the disease. Using our previously developed STEAP1 CAR T cell therapy, we evaluated preclinical efficacy across multiple orthotopic intratibial and disseminated EwS models. To overcome the immunosuppressive nature of EwS and enhance the therapeutic potential of STEAP1 CAR T cells, we armored them to constitutively express mature interleukin-18 (IL-18). IL-18 is a pro-inflammatory cytokine known to amplify T cell cytotoxicity by inducing interferon-gamma (IFN-γ) secretion from T cells and NK cells and enhancing granule-mediated killing and Fas ligand expression^29^. While systemic administration of recombinant IL-18 has demonstrated safety in clinical trials, its efficacy as monotherapy has been limited^30^. We show that IL-18 armored STEAP1 CAR T cells elicit robust antitumor activity in preclinical EwS models, reducing the cell dose needed to achieve therapeutic responses. Together, our findings provide compelling evidence that IL-18 armored STEAP1 CAR T cells can potentiate durable antitumor responses, supporting the rationale for future clinical translation to treat relapsed/refractory EwS.

## RESULTS

### EWS::FLI1 fusion drives STEAP1 expression in Ewing sarcoma

We first independently confirmed STEAP1 expression in EwS and its regulation by the oncogenic EWS::FLI1 fusion protein. Immunohistochemical (IHC) staining for STEAP1 was performed on 34 archived clinical EwS tumor specimens at the Department of Pathology, Duke University Medical Center. We observed widespread plasma membrane localization of STEAP1 across the cohort (**Figure 1A**), consistent with prior studies^21,31^. A research pathologist scored STEAP1 expression in each sample using a semi-quantitative H-score based on staining intensity and percentage of positive tumor cells. 33 out of 34 samples exhibited STEAP1 expression above the threshold (H-score >30), while one sample showed lower expression with an H-score of 20 (**Figure 1A–B**). To confirm the previously reported regulation of STEAP1 by EWS::FLI1^21^, we interrogated the publicly available Ewing Sarcoma Cell Line Atlas (ESCLA) dataset (GSE176339)^32^, in which 18 human EwS cell lines were stably transduced with a doxycycline (dox)-inducible short hairpin RNA (shRNA) targeting the EWSR1-ETS fusion transcript (via FLI1) followed by RNA sequencing^33^. Upon dox-induced knockdown of EWSR1-ETS, there was a significant downregulation of STEAP1 transcript levels across all cell lines (**Figure 1C**), consistent with transcriptional regulation by EWS::FLI1. To investigate regulation at the protein level, we examined publicly available proteomic data (PXD018937) from experiments analyzing time dependent protein changes upon targeted degradation of an FKBP12^F36V^-tagged EWS::FLI1 fusion protein using the dTAG system^34^. Degradation of EWS::FLI1 was observed within two hours of dTAG^V^-1 treatment and protein loss was maintained over twenty-four hours. Notably, STEAP1 protein levels followed similar kinetics and decreased significantly within two hours of EWS/FLI1 degradation (**Figure 1D**), further supporting that the fusion oncoprotein directly regulates STEAP1 expression with similar effects at both the transcript and protein levels in EwS.

**Figure 1.**
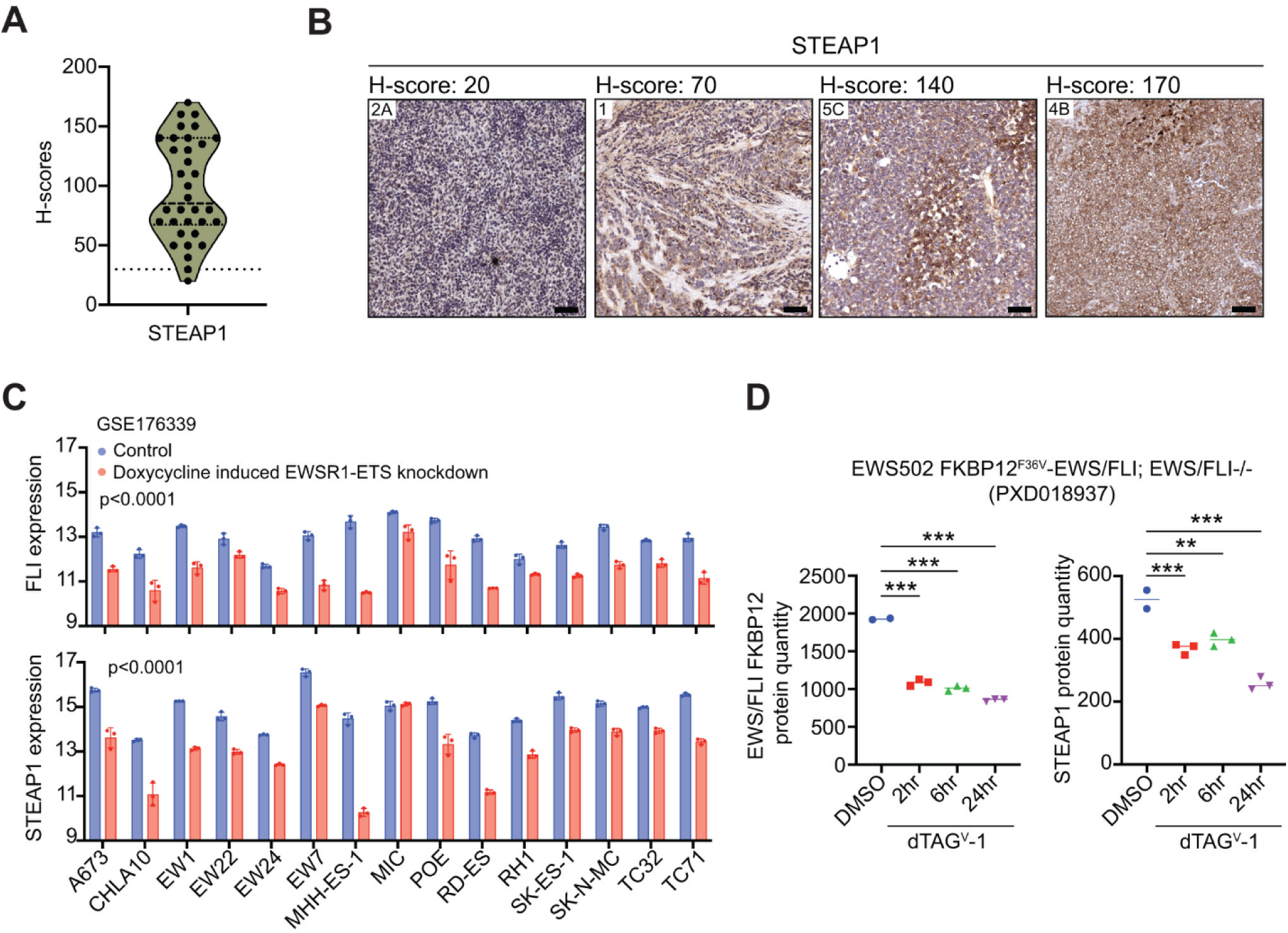
STEAP1 expression in EwS and its regulation by EWS::FLI1. (A) Quantification of STEAP1 protein expression in 34 archived clinical EwS tumor samples by IHC. H-scores were calculated as (% positive tumor cells × staining intensity [scale 0–2]). (B) Representative IHC images showing membranous STEAP1 staining in tumors with varying H-scores: 20 (STEAP1-negative), 70, 140, and 170 (STEAP1-positive). (C) Relative FLI1 (top) and STEAP1 (bottom) mRNA expression in EwS cell lines from the Ewing Sarcoma Cell Line Atlas (GSE176339) following doxycycline-inducible shRNA-mediated FLI1 knockdown. P-values were calculated using two-way ANOVA with Šídák’s multiple comparisons test; p < 0.0001. Data are presented as mean ± SD. (D) Proteomic quantification of EWS::FLI1 (left) and STEAP1 (right) in EWS502 cells treated with DMSO or dTAG^V^-1 to induce targeted degradation of FKBP12^F36V^-EWS::FLI1. P-values were calculated using one-way ANOVA with Dunnett’s multiple comparisons test; **p<0.001, ***p<0.0001. Data are shown as scatter plots with median.

### Targeting EWS/FLI-induced STEAP1 with STEAP1 CAR T cell therapy in Ewing sarcoma

The EWSR1::FLI1 fusion in EwS is commonly classified into molecular subtypes based on the genomic breakpoint locations. The most prevalent is type I (∼60%) defined by a fusion of EWSR1 exon 7 to FLI1 exon 6 and type II (∼25%) defined by fusion of EWSR1 exon 7 to FLI1 exon 5. A less common but distinct type III fusion involves EWSR1 exon 7 fused to FLI1 exon 8, resulting in a longer chimeric protein^35,36^. In approximately 5–10% of cases, EWSR1 fuses with other ETS family members such as ERG, ETV1, E1AF, and FEV, forming variant EWSR1::ETS fusions^37,38^. These distinct fusions have been shown to differentially influence epigenetic and transcriptional heterogeneity,^33^ biologic features that may also have clinical implications. ^39^.

To examine how EWSR1-FLI1 fusion subtypes influence STEAP1 expression, we analyzed a panel of human EwS cell lines representing distinct fusion types: type I (A673, TC32, TC71, SK-N-MC, CHLA10); type II (RD-ES, SK-ES-1); and type III (A4573). Flow cytometry showed STEAP1 surface expression across most EwS lines, irrespective of fusion subtype. TC-71, SK-N-MC and A4573 showed the highest relative STEAP1 expression while RD-ES and SK-ES-1 demonstrated the lowest (**Figure 2A**). The prostate cancer cell line DU145, which lacks STEAP1 expression, served as a negative control. Building on these findings, we evaluated the activation and cytotoxicity of our STEAP1 CAR T cell therapy^28^ (referred to as STEAP1-BBζ) across human EwS cell lines. This CAR incorporates a fully human single-chain variable fragment derived from the antibody-drug conjugate DSTP3086S, an IgG4 hinge CH2-CH3 long spacer containing the 4/2-NQ mutation^40^, a 4-1BB co-stimulatory domain, a CD3ζ activation domain, and a truncated epidermal growth factor receptor (EGFRt) as a transduction marker. Lentiviral vectors encoding the STEAP1-BBζ CAR were used to transduce CD4⁺ and CD8⁺ T cells, which were enriched from peripheral blood mononuclear cells (PBMCs) obtained from healthy human donors via leukapheresis. Following expansion, CAR T cells were immunophenotyped (**Supplementary Figure 1**) and reconstituted into a defined, normal CD4/CD8 ratio to enable standardized functional assessment. To assess CAR T cell activation, we performed immunologic co-culture assays between STEAP1-BBζ CAR T cells and a panel of EwS cell lines with measurement of IFN-γ secretion at 48 hours. We observed a robust increase in IFN-γ production in all EwS cell lines tested but not in the DU145 cell line (**Figure 2B**), indicating broad activation of STEAP1-BBζ CAR T cells when subjected to co-culture with EwS models. To investigate cytotoxic killing activity, we conducted time-lapse live-cell imaging of immunologic co-cultures using three EwS target cell lines: SK-ES-1 and RD-ES (relatively low STEAP1) and SK-N-MC (relatively high STEAP1). Each was co-cultured with either STEAP1-BBζ CAR T cells or untransduced T cells as a control. Across all three cell lines, STEAP1-BBζ CAR T cells induced robust and sustained cytolysis, demonstrating potent target cell killing regardless of the degree of STEAP1 expression (**Figure 2C-E**).

**Figure 2.**
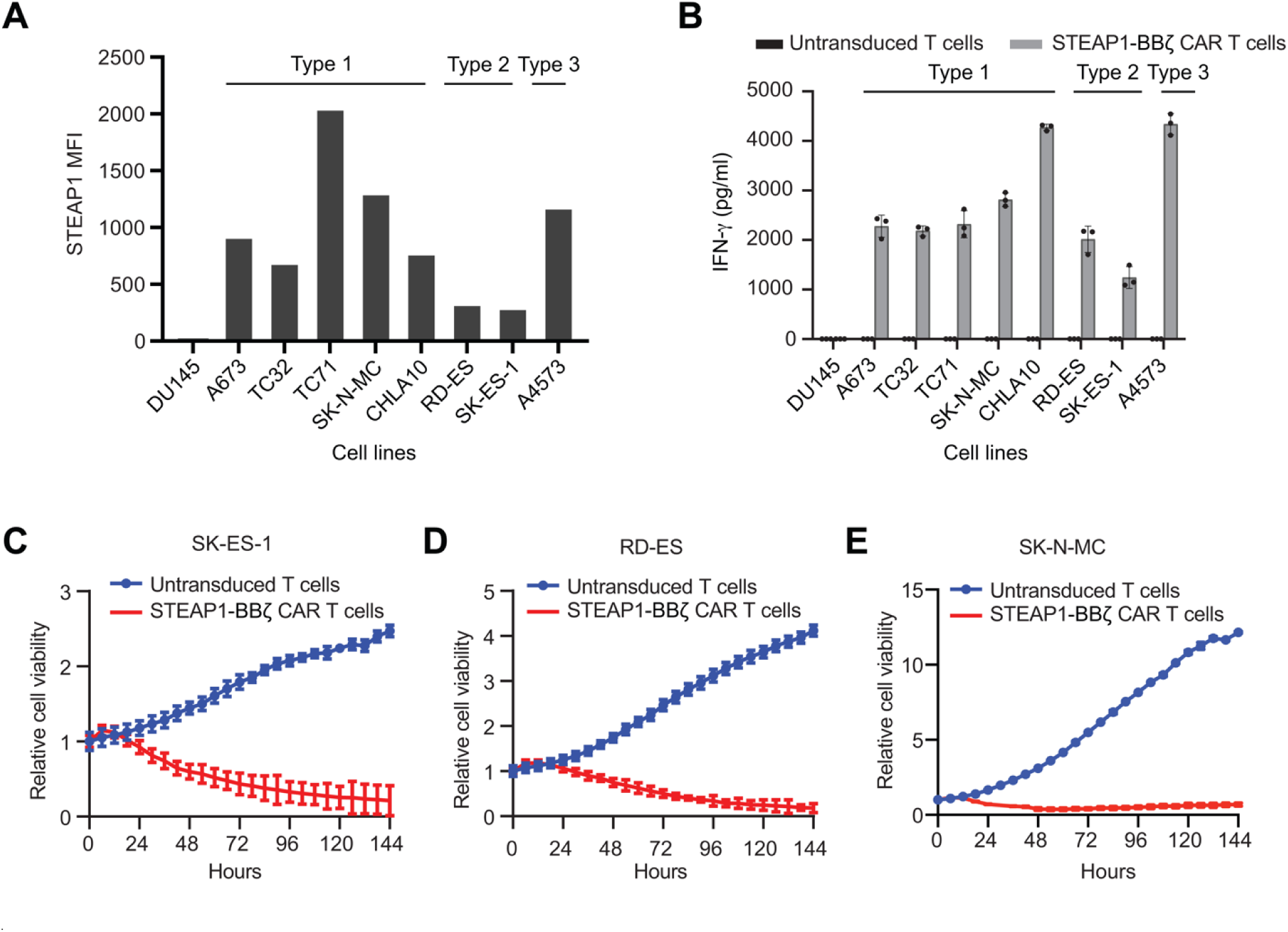
Activation and cytotoxicity of STEAP1-BB**ζ** CAR T cells against *in vitro* models of EwS. (A) Flow cytometry analysis of STEAP1 surface expression in EwS cell lines including type I, type II, and type III EWS::FLI1 fusions. Mean fluorescence intensity (MFI) values are shown. DU145 prostate cancer cells were used as a negative control. (B) IFN-γ ELISA results at 48 hours from immunologic co-cultures of either untransduced T cells or STEAP1-BBζ CAR T cells with different EwS cell lines at a 1:1 ratio. (C-E) Relative cell viability of (C) SK-ES-1 (D) RD-ES and (E) SK-N-MC cells over time measured by fluorescence live-cell imaging upon immunologic co-culture with either untransduced T cells or STEAP1-BBζ CAR T cells at a 1:1 ratio.

### STEAP1-BB**ζ** CAR T cells exhibit potent antitumor activity in orthotopic intratibial models of Ewing sarcoma

To assess the *in vivo* efficacy of STEAP1-BBζ CAR T cells in a clinically relevant context, we established orthotopic xenograft models using firefly luciferase (fLuc)-expressing SK-ES-1 and RD-ES cell lines. These EwS cell lines were engrafted into the tibia of immunodeficient NSG mice, recapitulating the primary bone site commonly affected in human EwS. Tumor progression and metastatic spread were monitored via serial bioluminescence imaging (BLI) starting one to two weeks post-engraftment. Upon confirmation of tumor establishment, mice received a single intravenous infusion of either 5 × 10^6^ STEAP1-BBζ CAR T cells or untransduced T cells as a control (**Figure 3A**).

**Figure 3.**
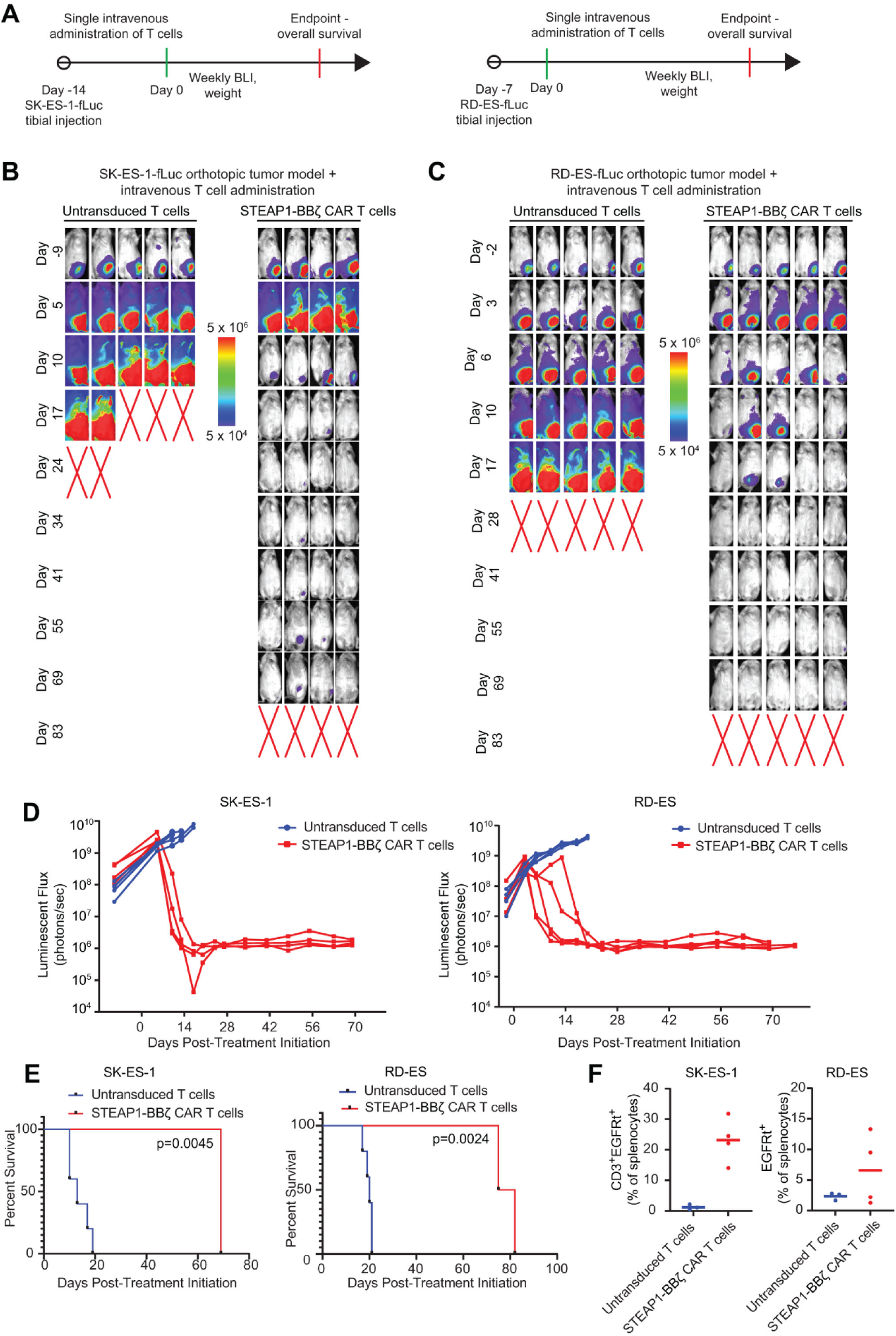
*In vivo* antitumor activity of STEAP1-BB**ζ** CAR T cells in orthotopic EwS models. (A) Schematic of tumor challenge experiments for SK-ES-1(left) and RD-ES (right) orthotropic (intratibial) mice models. fLuc=firefly luciferase, BLI=bioluminescence imaging. (B) Serial BLI of NSG mice orthotopically engrafted with SK-ES-1-fLuc^+^ cells and treated with a single intravenous injection of 5L×L10^6^ untransduced T cells (n=5) or STEAP1-BBζ CAR T cells (n=4) on day 0. Red X denotes deceased mice. Radiance scale is shown. (C) Serial BLI of NSG mice orthotopically engrafted with RD-ES-fLuc^+^ cells and treated with a single intravenous injection of 5L×L10^6^ untransduced T cells (n=5) or STEAP1-BBζ CAR T cells (n=5) on day 0. Red X denotes deceased mice. Radiance scale is shown. (D) Plot showing the quantification of total flux over time from live BLI of each mouse in (B) and (C). (E) Kaplan–Meier survival curves of mice in (B) and (C) with statistical significance determined by log-rank (Mantel-Cox) test. (F) Quantification of CD3^+^EGFRt^+^ STEAP1-BBζ CAR T cells by flow cytometry from splenocytes of mice treated with untransduced or STEAP1-BBζ CAR T cells at the end of the study on day 83. Bars represent mean.

Treatment with STEAP1-BBζ CAR T cells resulted in a marked reduction in tumor burden across both orthotopic EwS models (**Figure 3B–C**). Serial BLI imaging revealed progressive tumor growth in mice receiving untransduced T cells whereas mice treated with STEAP1-BBζ CAR T cells exhibited complete responses (**Figure 3D**). Importantly, STEAP1-BBζ CAR T cell therapy was associated with prolonged survival relative to treatment with untransduced T cells with a median survival of 83 vs. 17 days in the SK-ES-1 model (p = 0.0045) and from 83 vs. 28 days in the RD-ES model (p = 0.0024) as determined by log-rank test (**Figure 3E**). Mice in the treatment group were euthanized at day 83 for end point analysis. No significant changes in body weight were observed in mice treated with STEAP1-BBζ CAR T cells in either model (**Supplementary Figure 2**).

To assess persistence of STEAP1-BBζ CAR T cells, mice were euthanized at the study end point on day 83, splenocytes were harvested and flow cytometry was performed to quantify CAR^+^ T cells based on co-expression of CD3 and the EGFRt transduction marker. We observed robust persistence of CD3⁺ EGFRt⁺ STEAP1-BBζ CAR T cells in treated mice across both models (**Figure 3F**). These findings demonstrate that STEAP1-BBζ CAR T cell therapy results in potent and durable antitumor responses in both orthotopic EwS models. Further, they underscore the clinical potential of this therapeutic approach by validating the antitumor activity of STEAP1-BBζ CAR T cells within the bone microenvironment and their persistence.

### Dose escalation defines minimum effective threshold for STEAP1-BB**ζ** CAR T cell therapy

To inform clinically relevant dosing strategies and address current limitations in CAR T cell manufacturing and scalability^41,42^, we conducted a dose escalation study to evaluate the minimum effective dose of STEAP1-BBζ CAR T cells *in vivo*. For this analysis, we employed a disseminated EwS model using the fLuc-expressing SK-ES-1 cell line administered via tail vein injection in NSG mice to approximate metastatic EwS. Twenty days post-engraftment after confirmation of metastatic colonization by BLI, mice were randomized into four groups and treated with a single intravenous dose of either 5 × 10^6^, 10^6^, or 2 × 10^5^ STEAP1-BBζ CAR T cells or 5 × 10^6^ untransduced T cells as a control (**Figure 4A**).

**Figure 4.**
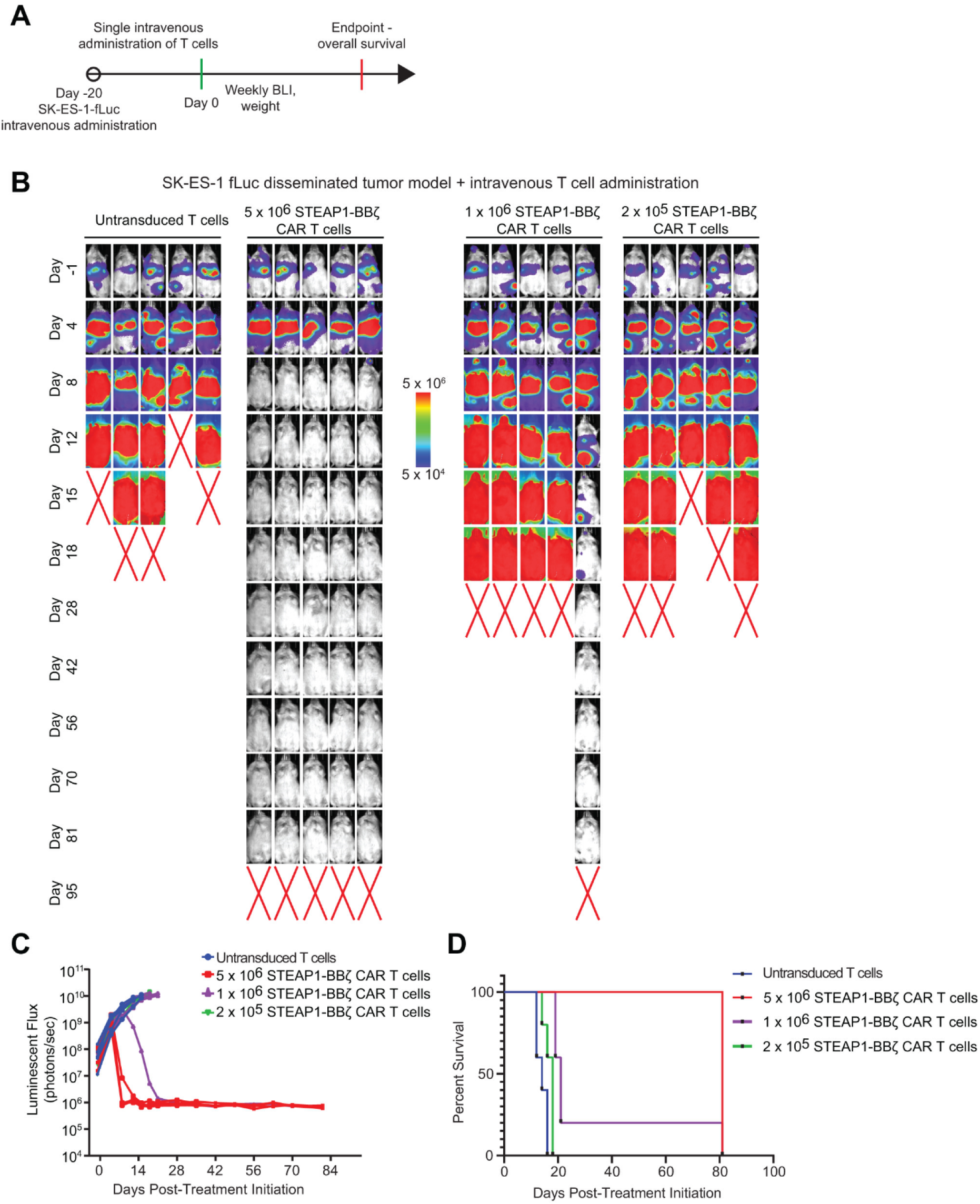
Dose-dependent therapeutic response of STEAP1-BB**ζ** CAR T cells in disseminated EwS models. (A) Schematic of tumor challenge experiments for disseminated SK-ES-1-fluc^+^ models. (B) Serial BLI of NSG mice engrafted by tail vein injection with SK-ES-1-fLuc^+^ cells and treated with a single intravenous injection of either 5 ×L10^6^ untransduced T cells (n=5) or 5L×L10^6^, 10^6^, and 2L×L10^5^ STEAP1-BBζ CAR T cells (n=5 each) a on day 0. Red X denotes deceased mice. Radiance scale is shown. (C) Plot showing the quantification of total flux over time from live BLI of each mouse in (B). (D) Kaplan–Meier survival curves of mice in (B) with statistical significance determined by log-rank (Mantel-Cox) test.

Serial BLI revealed a dose-dependent antitumor response with mice treated with the highest dose of 5 × 10^6^ STEAP1-BBζ CAR T cells demonstrating complete responses in all five animals (**Figure 4B–C**). At the 10^6^-cell dose, only one of five mice exhibited a complete response while no therapeutic benefit was observed at the 2 × 10^5^ cell dose. Survival analysis showed a significant extension in median survival associated with the 5 × 10^6^ STEAP1-BBζ CAR T cell dose (85 vs. 17 days, p < 0.0001), whereas the 10^6^-cell dose group showed partial survival benefit and the 2 × 10^5^ cell dose group displayed survival outcomes similar to the control untransduced T cell group (**Figure 4D**). These results highlight a potential dose threshold effect for therapeutic efficacy.

No significant changes in body weight were observed across any of the treatment groups, indicating that CAR T cell administration was well tolerated (**Supplementary Figure 3A**). To investigate whether treatment failure at lower cell doses may have been a consequence of antigen escape, as previously seen in studies in disseminated prostate cancer models^28^, we performed IHC analysis for STEAP1 expression on residual tumor tissues collected at the study endpoint. All residual tumors retained high STEAP1 expression regardless of treatment dose, ruling out acute antigen loss as a mechanism of resistance (**Supplementary Figure 3B**). These findings highlight a critical dose threshold required for effective antitumor activity and underscore the need for further optimization strategies to further empower STEAP1-BBζ CAR T cells to achieve durable responses at lower, clinically scalable doses.

### IL-18 armoring enhances STEAP1-BB**ζ** CAR T cells therapeutic efficacy at lower cell dose

To overcome the reduced efficacy of STEAP1-BBζ CAR T cells at lower cell doses and to promote more potent antitumor responses, we engineered a cytokine-armored CAR construct that constitutively expresses mature human IL-18 and subsequently generated STEAP1-BBζ-hIL18 CAR T cells (**Figure 5A**). To assess the functional advantage of IL-18 armoring, we performed an *in vitro* tumor rechallenge assay using the SK-ES-1 cell line. Tumor cells were co-cultured with either STEAP1-BBζ or STEAP1-BBζ-hIL18 CAR T cells and subjected to serial rechallenges every 48 hours at defined effector-to-target (E:T) ratios. Under increasingly stringent conditions using E:T ratios down to 1:8, we observed a marked decline in the cytotoxic activity of unarmored STEAP1-BBζ CAR T cells (**Figure 5B**). In contrast, STEAP1-BBζ-hIL18 CAR T cells maintained superior tumor-killing capacity across multiple rounds of tumor rechallenge. Enzyme-linked immunosorbent assay (ELISA) analysis of culture supernatants collected at both no-rechallenge and rechallenged conditions showed significantly elevated levels of both human IL-18 (**Figure 5C**) and IFN-γ (**Figure 5D**) associated with the STEAP1-BBζ-hIL18 CAR T cell group, indicating enhanced effector cytokine production upon tumor cell engagement.

**Figure 5.**
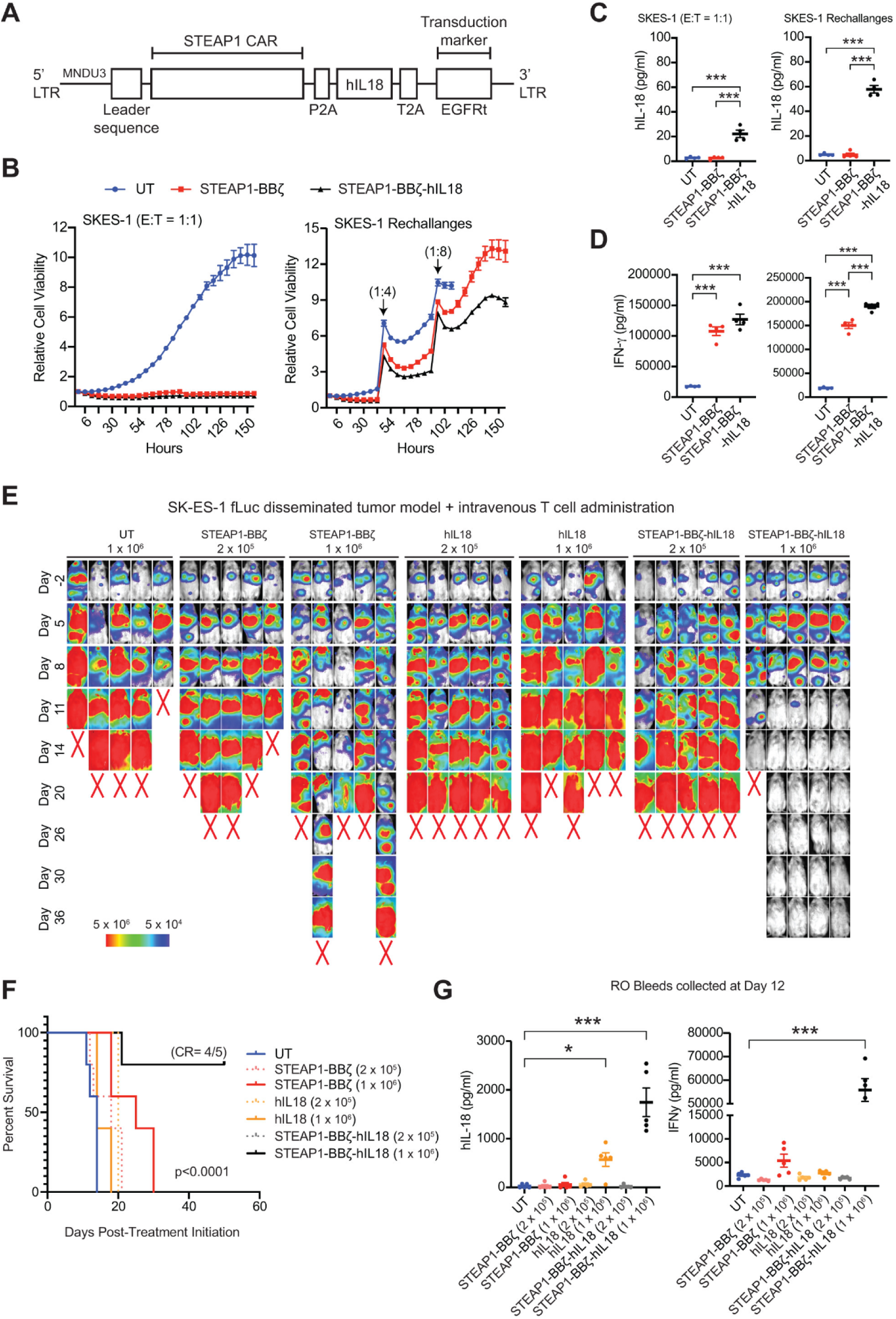
IL-18 armored STEAP1-BB**ζ** CAR T cells exhibit potent antitumor activity at a lower cell dose. (A) Schematic representation of the fourth-generation lentiviral STEAP1-BBζ CAR construct armored with human IL-18 (hIL18). The construct includes the previously defined STEAP1-targeting CAR domains, with hIL-18 flanked by P2A self-cleaving peptide sequences. LTR=long terminal repeat, EGFRt=truncated epidermal growth factor receptor. (B) Relative viability of SK-ES-1 cells over time, assessed by fluorescence-based live-cell imaging during immunologic co-culture with either untransduced T cells (UT), unarmored STEAP1-BBζ CAR T cells, or hIL-18–armored STEAP1-BBζ-hIL18 (STEAP1-BBζ-hIL18) CAR T cells at an effector-to-target (E: T) ratio of 1:1, including serial tumor re-challenges at specific E:T ratios every 48 hours. (C) Human IL-18 ELISA results from immunologic co-culture supernatants in (B) after 48 hours. P-values were calculated using two-way ANOVA with Šídák’s multiple comparisons test; ***pL<L0.0001. Data are shown as scatter plots with median. (D) Human IFN-γ ELISA results from immunologic co-culture supernatants in (B) after 48 hours. P-values were calculated using two-way ANOVA with Šídák’s multiple comparisons test; ***pL<L0.0001. Data are shown as scatter plots with median. (E) Serial BLI of NSG mice engrafted by tail vein injection with SK-ES-1-fLuc^+^ cells and treated with a single intravenous injection of either 5 ×L10^6^ untransduced T cells (n=5) or 5L×L10^6^,L10^6^, and 2L×L10^5^ STEAP1-BBζ CAR T cells (n=5 each) on day 0. Red X denotes deceased mice. Radiance scale is shown. (F) Kaplan–Meier survival curves of mice in (B) with statistical significance determined by log-rank (Mantel-Cox) test. (G) Human IL-18 (left) and IFN-γ (right) ELISA results from retroorbital (RO) bleeds collected at day 12 from mice in (E). P-values were calculated using two-way ANOVA with Šídák’s multiple comparisons test; *pL<L0.01, ***pL<L0.0001. Data are shown as scatter plots with median.

To validate the enhanced therapeutic potential of IL-18 armoring observed *in vitro*, we next directly compared the *in vivo* antitumor efficacy of STEAP1-BBζ and STEAP1-BBζ-hIL18 CAR T cells in the SK-ES-1 disseminated EwS model. After confirmation of metastatic colonization, mice were randomized into seven groups and treated with a single dose of either 2 × 10^5^ or 10^6^ hIL-18 T cells, STEAP1-BBζ CAR T cells, or STEAP1-BBζ-hIL18 CAR T cells or 10^6^ untransduced T cells as a control. Flow cytometry was performed to confirm expression of STEAP1-BBζ CAR or STEAP1-BBζ-hIL18 CAR in the cell products (**Supplementary Figure 4A**).

Serial BLI revealed a marked reduction in tumor burden exclusively in mice treated with 10^6^ STEAP1-BBζ-hIL18 CAR T cells (**Figure 5E, Supplementary Figure 4B**), where four of five mice achieved complete responses (**Figure 5F**). A transient 10% reduction in body weight was observed in this group which stabilized over time (**Supplementary Figure 4C**). To investigate systemic cytokine responses, retroorbital blood was collected from all groups. Mice treated with 10^6^ STEAP1-BBζ-hIL18 CAR T cells exhibited significantly elevated circulating levels of human IL-18 and IFN-γ at day 12, coinciding with peak therapeutic response. Together, these findings demonstrate that IL-18 armoring enhances the antitumor potency of STEAP1-BBζ CAR T cells *in vivo*, enabling effective tumor clearance at lower cell doses without gross toxicity.

## DISCUSSION

EWS-FLI1 is the central oncogenic driver of EwS, acting as an aberrant transcription factor that reshapes the epigenetic and transcriptional landscape of tumor cells^2,43^. Despite its pivotal role, EWS-FLI1 remains challenging to target directly; however, its downstream target STEAP1 has emerged as a promising immunotherapeutic candidate^31,44,45^. STEAP1 is transcriptionally upregulated through the cooperative interaction of EWS-FLI1 and its downstream effector NKX2.2, which binds to adjacent regulatory elements at the STEAP1 promoter to enhance its expression^44^. Importantly, STEAP1 displays minimal expression in normal adult tissues, further supporting its potential as a tumor-selective target that may be associated with a favorable therapeutic index. In this study, we demonstrate that STEAP1 is widely expressed in ∼97% of EwS tumors and across multiple EWS::FLI1 fusion-driven subtypes of human EwS cell lines, consistent with prior reports showing STEAP1 expression across EwS harboring alternative ETS fusions^21^. We further validate that STEAP1 expression is directly regulated by the EWS-FLI1 fusion protein at both the RNA and protein levels, as perturbation of EWS-FLI1 results in a marked decline in STEAP1 expression.

STEAP1 was recently established as a clinically validated cancer target given that the STEAP1×CD3 TCE xaluritamig demonstrated significant clinical activity with tolerable toxicities in a phase I clinical trial for mCRPC^27^. Recently, a multitude of STEAP1-targeted therapeutic agents for cancer have also entered into preclinical and clinical development. CAR T cell therapies may provide specific advantages as they may provide long-term immune surveillance and are readily amenable to engineering strategies to overcome the hostile, immunosuppressive TME of solid tumors. We previously engineered STEAP1 CAR T cells to target prostate cancer with low antigen density and demonstrated potent antitumor activity across several preclinical models^28^. In this study, we show that STEAP1 CAR T cells also induce tumor eradication of both orthotopic and disseminated human EwS models. Even with dose de-escalation, we found that STEAP1 CAR T cells could induce complete responses in a subset of mice without signs of antigen escape. These findings contrast with the recurrent STEAP1 loss previously observed in progressive or relapsed tumors after STEAP1 CAR T cell therapy in prostate cancer models^28^. We suspect that this could be a consequence of STEAP1 expression being inexorably coupled with the activity of the oncogenic driver and tumor dependency EWS-FLI1 in EwS.

Armoring CAR T cell therapies to express recombinant cytokines has emerged as a leading strategy to enhance the potency of CAR T cell therapies by improving cytotoxic effector function, enhancing expansion and persistence, and promoting endogenous antitumor immunity. Among these, interleukins such as IL-12^46,47^, IL-15^48^, and IL-18^49^ have shown profound preclinical efficacy in augmenting the antitumor activity of CAR T cells. IL-18 is a proinflammatory cytokine that enhances T and NK cell cytotoxicity and promotes IFN-γ production. When mature IL-18 was expressed from CAR T cells targeting DLL3^50^ or CD19^49^, it was found to remodel the immunosuppressive TME by recruiting and activating innate and adaptive immune cells. IL-18 has also been shown to promote macrophage polarization toward an M1 phenotype, activate dendritic cells, and improve antigen presentation, thereby enhancing endogenous T cell priming and antitumor immunity^50–52^. These immunomodulatory effects have translated into early clinical success based on the results of a clinical trial evaluating IL-18-secreting CD19 CAR T cells (huCART19-IL18)^49^ in patients with relapsed or refractory B-cell lymphoma after commercial CD119 CAR T cell therapy (NCT04684563) which showed a response rate of ∼81%, durable clinical benefit, and a favorable safety profile. Notably, the huCART19-IL18 study demonstrated that despite secretion of bioactive IL-18, systemic immune homeostasis was preserved, likely due to the buffering capacity of endogenous IL-18 binding protein (IL-18BP) that regulates free circulating IL-18^53^. The positive correlation between IL-18:IL-18BP complex levels and CAR T cell expansion suggests IL-18 acts primarily within the TME without disrupting systemic cytokine balance. In line with these findings, our preclinical study of IL-18 armored STEAP1 CAR T cells in a disseminated EwS model showed complete tumor regression in ∼80% of mice treated at a dose of 10^6^ CAR T cells with no gross toxicities observed. Importantly, the same cell dose of unarmored STEAP1 CAR T cells was tested in parallel and incapable of eliciting complete responses in any of the mice. While limited by the lack of syngeneic mouse EwS models available for use in immunocompetent mice, these results still underscore the potential of IL-18 armoring to further enhance the therapeutic potency of STEAP1 CAR T cell therapy in EwS.

Despite the transformative success of CAR T cell therapy in hematologic malignancies, its efficacy in solid tumors remains limited with no clinically approved therapies to date. However, select early-stage clinical trials have reported on the promising activity and safety of CAR T cell therapies in childhood cancers. GD2-targeted CAR T cell therapy has demonstrated encouraging results in H3K27M-mutant diffuse midline glioma ^54,55^ and neuroblastoma^56^, including a first-generation GD2 CAR T cell therapy that induced long-term remission lasting over 18 years in a neuroblastoma patient^57^. More advanced GD2 CAR designs such as GD2-CART01 incorporating CD28, OX40, CD3ζ, and an inducible caspase 9 (iC9) safety switch have shown objective response rates of 63% and durable complete remissions in relapsed/refractory neuroblastoma^56^. While STEAP1 CAR T cell therapy is currently being evaluated in a phase I/II trial for mCRPC (NCT06236139), we plan to initiate a phase I trial of STEAP1 CAR T cell therapy for relapsed/refractory EwS based on the compelling results of this study. In addition, early planning of a future clinical trial of IL-18 armored STEAP1 CAR T cell therapy has also commenced.

## METHODS

### Cell lines

All Ewing sarcoma cell lines used in this study—A673, TC32, TC71, SK-N-MC, CHLA10, RD-ES, SK-ES-1, and A4573—were kindly provided by Dr. Elizabeth Lawlor (Seattle Children’s Research Institute, Seattle, WA, USA). The prostate cancer cell lines DU145 (HTB-81), 22Rv1 (CRL-2505), C4-2B (CRL-3315), and the HEK293T cell line (CRL-3216) were obtained from the American Type Culture Collection (ATCC). RD-ES, TC32, A4573, 22Rv1, C4-2B, and DU145 cell lines were cultured in RPMI 1640 medium supplemented with 10% fetal bovine serum (FBS), 100 U/mL penicillin, 100 µg/mL streptomycin, and 4 mmol/L GlutaMAX (Thermo Fisher Scientific). CHLA10 and TC71 cells were cultured in Iscove’s Modified Dulbecco’s Medium (IMDM) supplemented with 20% FBS, 4 mmol/L L-glutamine, and 1× ITS (5 µg/mL insulin, 5 µg/mL transferrin, 5 ng/mL selenous acid). A673 and HEK293T cells were cultured in Dulbecco’s Modified Eagle’s Medium (DMEM) supplemented with 10% FBS, 100 U/mL penicillin, 100 µg/mL streptomycin, and 4 mmol/L GlutaMAX. SK-N-MC cells were cultured in Eagle’s Minimum Essential Medium (EMEM) with 10% FBS, 100 U/mL penicillin, 100 µg/mL streptomycin, and 4 mmol/L GlutaMAX.

### Animal studies

All mouse studies were conducted at the Fred Hutchinson Cancer Center in accordance with protocols approved by the Institutional Animal Care and Use Committee (IACUC) and in compliance with the regulations of Comparative Medicine.

For intra-tibial and disseminated tumor models, six- to eight-week-old male NSG (NOD-SCID IL2Rγ^null) immunodeficient mice were obtained from The Jackson Laboratory and used for all in vivo experiments. In the intra-tibial model, 1 × 10^6^ SK-ES-1 or RD-ES tumor cells were resuspended in 100 µL of ice-cold Matrigel Matrix (Corning) and injected directly into the tibia. For the disseminated model, 1 × 10^6^ SK-ES-1 tumor cells were resuspended in 100 µL of PBS and injected intravenously via the tail vein. Tumor burden was monitored longitudinally using bioluminescence imaging (BLI) on the IVIS Spectrum system (PerkinElmer) following intraperitoneal injection of XenoLight D-luciferin (PerkinElmer). Upon establishment of metastatic colonization, mice received a tail vein injection of 100 µL PBS containing a defined number of human STEAP1-BBζ, STEAP1-BBζ-IL18, or untransduced CAR T cells.

At humane endpoints, mice were euthanized, and metastatic tumors and spleens were harvested. Tumors were formalin-fixed and paraffin-embedded for histopathological analysis. Splenocytes were isolated for flow cytometric analysis to evaluate the peripheral persistence of CAR T cells.

### STEAP1 immunohistochemistry

Formalin-fixed, paraffin-embedded (FFPE) tissue sections from Ewing sarcoma patients were obtained from Dr. Jiaoti Huang, Department of Pathology, Duke University Medical Center. STEAP1 IHC on EWS patient tumor samples and mice tumor tissues was performed as previously described^28^. Briefly, FFPE sections were deparaffinized in xylene, rehydrated through a graded ethanol series (100%, 95%, and 75%), and washed in Tris-buffered saline with 0.1% Tween-20 (TBST). Antigen retrieval was conducted using Citrate-Based Antigen Unmasking Solution (Vector Laboratories) in a pressure cooker at 95°C for 30 minutes. Dual Endogenous Enzyme-Blocking Reagent (Agilent Technologies) was applied for 10 minutes, followed by TBST washes. Slides were incubated with a primary rabbit anti-STEAP1 antibody (LS Bio, LS-C291740, 1:500) at 37°C for 1 hour in a humidified chamber, washed, and then incubated with PowerVision Poly-HRP anti-rabbit secondary antibody (Leica Biosystems) for 30 minutes at 37°C. After additional washes, slides were developed with 3,3′-diaminobenzidine (DAB; Sigma-Aldrich) for 5 minutes, counterstained with hematoxylin (Dako, Agilent Technologies) for 1 minute, dehydrated through graded ethanol, cleared in xylene, and mounted using Permount Mounting Medium (Fisher Chemical). Whole-slide images were acquired using a Ventana DP200 slide scanner (Roche) and analyzed with QuPath software (version 0.2.3).

STEAP1 staining intensity on the EwS patient tissues were scored by scored by an experienced pathologist (E.S) in a blinded fashion, using a semi-quantitative scale an intensity score was provided based on staining intensity (0 = none, 1 = weak, 2 = moderate, 3 = strong) and percentage of positive tumor cells. H-score for each sample was calculated as the product of intensity and percentage of cells stained resulting in a minimum of 0 and maximum of 200. Samples with H-score of more than 30 were graded STEAP1 positive.

### Flow cytometry-based determination of surface expression of STEAP1

To evaluate cell surface expression of STEAP1 in Ewing sarcoma cell lines, 2.0 × 10^5^ cells per condition—unstained, secondary antibody only, and fully stained (primary plus secondary antibodies)—were harvested, counted, and transferred into 5 mL polystyrene FACS tubes (Corning). Cells were centrifuged at 1200 rpm for 3 minutes and resuspended in 100 µL of FACS buffer. For staining, the primary antibody vandortuzumab (anti-STEAP1; Creative Biolabs) was used at 60 µg/ml concentration per 100 µL FACS buffer and added only to the fully stained condition. All samples were incubated on ice for 30 minutes in the dark. After washing twice with PBS, the secondary antibody—an APC-conjugated rat anti-human IgG Fc antibody (BioLegend)—was added to both the stained and secondary-only samples at the same dilution (2 µL per 100 µL buffer), followed by a second 30-minute incubation on ice in the dark. Cells were washed twice with PBS and resuspended in 200 µL of FACS buffer for flow cytometric analysis. Data were acquired using a BD FACSCanto II (BD Biosciences) and analyzed with FlowJo software v10.8 (Tree Star).

### Chimeric antigen receptor expression plasmids

The human pCCl-c-MNDU3-STEAP1-4-1BBζ CAR used in this study was engineered with the DSTP3086S scFv, an IgG4 spacer, and a truncated EGFR (EGFRt) as a transduction marker, as previously described^28^. To generate IL-18–armored CAR constructs, a gBlock containing the human IL-18 gene (Integrated DNA Technologies) was synthesized with an IL-2 signal peptide and a self-cleaving P2A sequence and subsequently cloned into the STEAP1 CAR backbone. Resulting constructs encoding hIL-18 CAR and STEAP1-BBζ-hIL18 (armored) CARs were verified by whole plasmid sequencing (Plasmidsaurus).

### Lentivirus production

To generate STEAP1-BBζ and STEAP1-BBζ-hIL18 CAR lentivirus, HEK293T cells were thawed, cultured, and expanded in complete DMEM supplemented with 10% FBS, GlutaMAX, and Penicillin-Streptomycin. For lentiviral production, cells were seeded onto Culturex Poly-L-Lysine–coated plates (R&D Systems). Next day, cells were transfected with the CAR transfer plasmid (pCCl-c-MNDU3 STEAP1-BBζ or pCCl-c-MNDU3 STEAP1-BBζ-hIL18) along with the packaging plasmids pMDL, pVSVg, and pREV using FuGENE HD Transfection Reagent (Promega). At 18 hours post-transfection, sodium butyrate and HEPES were added to each plate at a final concentration of 20 mM per reagent. Eight hours later, media were aspirated, cells were washed with PBS and replaced with fresh complete DMEM supplemented with 20 mM HEPES. After 48 hours post-transfection, lentiviral supernatants were harvested and filtered through a 0.22 µm vacuum filter. The virus-containing supernatant was concentrated by ultracentrifugation in polypropylene conical tubes (Beckman Coulter) at 85,929 × g for 2 hours at 4°C using an Optima XE 90 ultracentrifuge (Beckman Coulter). Following centrifugation, excess supernatant was carefully aspirated, leaving a minimal residual volume to resuspend the lentiviral pellet. The concentrated lentivirus was aliquoted into cryovials and stored at −80°C until use.

### STEAP1 and IL-18 armored STEAP1 CAR T cell manufacturing

Peripheral blood mononuclear cells (PBMCs) were obtained from de-identified healthy donors under written informed consent through the Co-Operative Center for Excellence in Hematology at the Fred Hutchinson Cancer Center, with protocols approved by the Institutional Review Board. For CAR T cell generation, PBMCs were thawed, washed, and centrifuged at 600g for 5 minutes before resuspension in T cell culture media (TCM), consisting of AIM-V (Gibco) supplemented with 55 mM β-mercaptoethanol, human AB serum (Sigma), and GlutaMAX. CD8⁺ and CD4⁺ T cells were isolated using magnetic MicroBeads (Miltenyi Biotec) and activated separately using Dynabeads Human T-Activator CD3/CD28 (Thermo Fisher Scientific) following manufacturer instructions. CD8⁺ T cells were cultured in TCM supplemented with 50 U/mL IL-2 and 0.5 ng/mL IL-15, while CD4⁺ T cells were cultured in TCM supplemented with 5 ng/mL IL-7 and 0.5 ng/mL IL-15. Twenty-four hours after activation (Day 2), T cells were transduced with STEAP1-BBζ or STEAP1-BBζ-hIL18 lentivirus at an MOI of 10 in the presence of 10 µg/mL protamine sulfate, based on viral titers established in HEK293T cells. Forty-eight hours post-transduction, activation beads were removed, and T cells were phenotyped on Day 5. CAR T cells were expanded for 10–14 days for in vitro and in vivo studies, with cell counts taken every two days and cultures maintained at 1 x 10^6^ cells/mL in their respective CD4 or CD8 media.

### CAR T cell Immunophenotyping

Transduced T cells were counted using a hemocytometer, and 1.5 x 10^5^ T cells from each group, untransduced, STEAP1-BBζ CAR, and STEAP1-BBζ-hIL18 CAR were collected for immunophenotyping. CD4⁺ and CD8⁺ transduced T cells were stained using a cocktail of following antibodies: FITC-conjugated mouse anti-human CD8 (BD Biosciences, Cat. 555366), APC-Cy7–conjugated mouse anti-human CD3 (Invitrogen, Cat. 47-0036-42), and PE-conjugated mouse anti-human EGFR (Novus Biologicals, Cat. NBP2-75903PE). Staining was performed on ice for 30 minutes in the dark. Cells were then washed with PBS and analyzed on a BD FACSCanto II flow cytometer (BD Biosciences). Flow cytometry data were analyzed using FlowJo v10.8 (Treestar) to determine the transduction efficiency of the CAR T cells.

### In vitro co-culture and tumor rechallenge assay

To assess CAR T cell-mediated cytotoxicity, GFP-expressing EwS target cells were co-cultured with untransduced T cells, STEAP1-BBζ CAR T cells, or STEAP1-BBζ-hIL18 CAR T cells at varying effector-to-target (E:T) ratios in clear-bottom, black-walled 96-well plates (Corning). Effector cell numbers were normalized to CAR transduction efficiency. Plates were pre-coated with Poly-L-Lysine (R&D Systems) for 30 minutes, seeded with target cells, and incubated at 37°C for at least one hour to promote adherence before the addition of effector T cells. Co-cultures were maintained in a BioTek BioSpa 8 Automated Incubator (Agilent Technologies) and imaged every six hours over a six-day period using a BioTek Cytation 5 Cell Imaging Multi-Mode Reader. GFP^+^ target cell counts were quantified using Gen5 software (Agilent Technologies) based on the number of GFP-positive objects per unit area. To assess T cell activation, 25–50 µL of supernatant was collected at 48 hours and stored at –30°C for cytokine analysis. For re-challenge assays, tumor cells were reintroduced to the co-culture every 48 hours at 1:4 or 1:8 ratios (target: T-cell), with supernatant collected at each re-challenge to monitor changes in cytokine secretion.

### IFN-***γ*** and IL-18 ELISA

Cytokine concentrations in co-culture supernatants and retro-orbital (RO) blood samples were measured using sandwich ELISA. Human IFNγ levels were assessed using the BD Human IFNγ ELISA Set (BD Biosciences, Cat. 555142), and human total IL-18 was measured using the Human Total IL-18/IL-1F4 Quantikine ELISA Kit (R&D Systems, Cat. DL180), following the manufacturers’ protocols. Cell supernatants were diluted to 1:5 for IFNγ ELISA and 1:25 for hIL-18 ELISA in the assay diluent buffers. Absorbance was read at 450 nm with wavelength correction at 560 nm using a BioTek Cytation 3 Cell Imaging Multi-Mode Microplate Reader (Agilent Technologies).

## ACKNOWLEDGEMENTS

We are deeply grateful to the patients and their families, whose contributions made this research possible. We thank Dr. Jiaoto Huang for providing Ewing sarcoma patient tissue samples and Radhika Patel for help with IHC experiments. We acknowledge the Fred Hutch Cooperative Center of Excellence in Hematology (supported by U54 DK106829), as well as the Fred Hutch core facilities, including the Experimental Histopathology Shared Resource and the Flow Cytometry Shared Resource. We also thank the UCLA Health Jonsson Comprehensive Cancer Center Shared Resources supported by the NIH/NCI under award number P30 CA016042 and Dr. Owen Witte’s laboratory for access to the BD Canto flow cytometer. This work was supported by a Swim Across America/Seattle Cancer Care Alliance Award (J.K.L.), CureSearch for Children’s Cancer Acceleration Initiative Award (J.K.L.), a Prostate Cancer Foundation Young Investigator Award (V.B.), and a Department of Defense Prostate Cancer Research Program Early Investigator Research Award (HT9425-23-1-0089 to V.B.).

## AUTHOR CONTRIBUTIONS

V.B. Data curation, formal analysis, validation, investigation, visualization, methodology, writing-original draft, writing-review and editing. A.T. Data curation, formal analysis, validation, investigation, visualization, methodology, writing-review and editing. E.S. and M.C.H. H-scoring, Data curation, formal analysis, and writing-review and editing. T.C. Data curation and formal analysis. P.P.C. Data curation and formal analysis. K.L. Data curation and formal analysis. E.R.L. Data curation, providing EwS cell lines, formal analysis, and writing-review and editing. J.H. Data curation, providing EwS patient FFPE tissues, formal analysis, and writing-review and editing. B.N. Data curation, formal analysis, and writing-review and editing. J.K.L. Conceptualization, data curation, formal analysis, validation, investigation, visualization, methodology, writing-original draft, writing-review and editing.

## DISCLOSURES

J.K.L. is an inventor on international patent applications (PCT/US2023/062428) for STEAP1 CAR T cell therapy. J.K.L. holds equity in, serves as Chief Medical Officer for, and has received research funding from PromiCell, Inc. J.K.L. is also a consultant for Lyell Immunopharma and Xilio Therapeutics. M.C.H. served as a paid consultant/received honoraria from Pfizer and Astra Zeneca and has received research funding from Merck, Novartis, Genentech and Bristol Myers Squibb. B.N. is an inventor on patent applications related to the dTAG system (WO/2017/024318, WO/2017/024319, WO/2018/148440, WO/2018/148443, and WO/2020/146250). The Nabet laboratory receives or has received research funding from Mitsubishi Tanabe Pharma America, Inc.

## SUPPLEMENTARY FIGURES

**Supplementary Figure 1.**
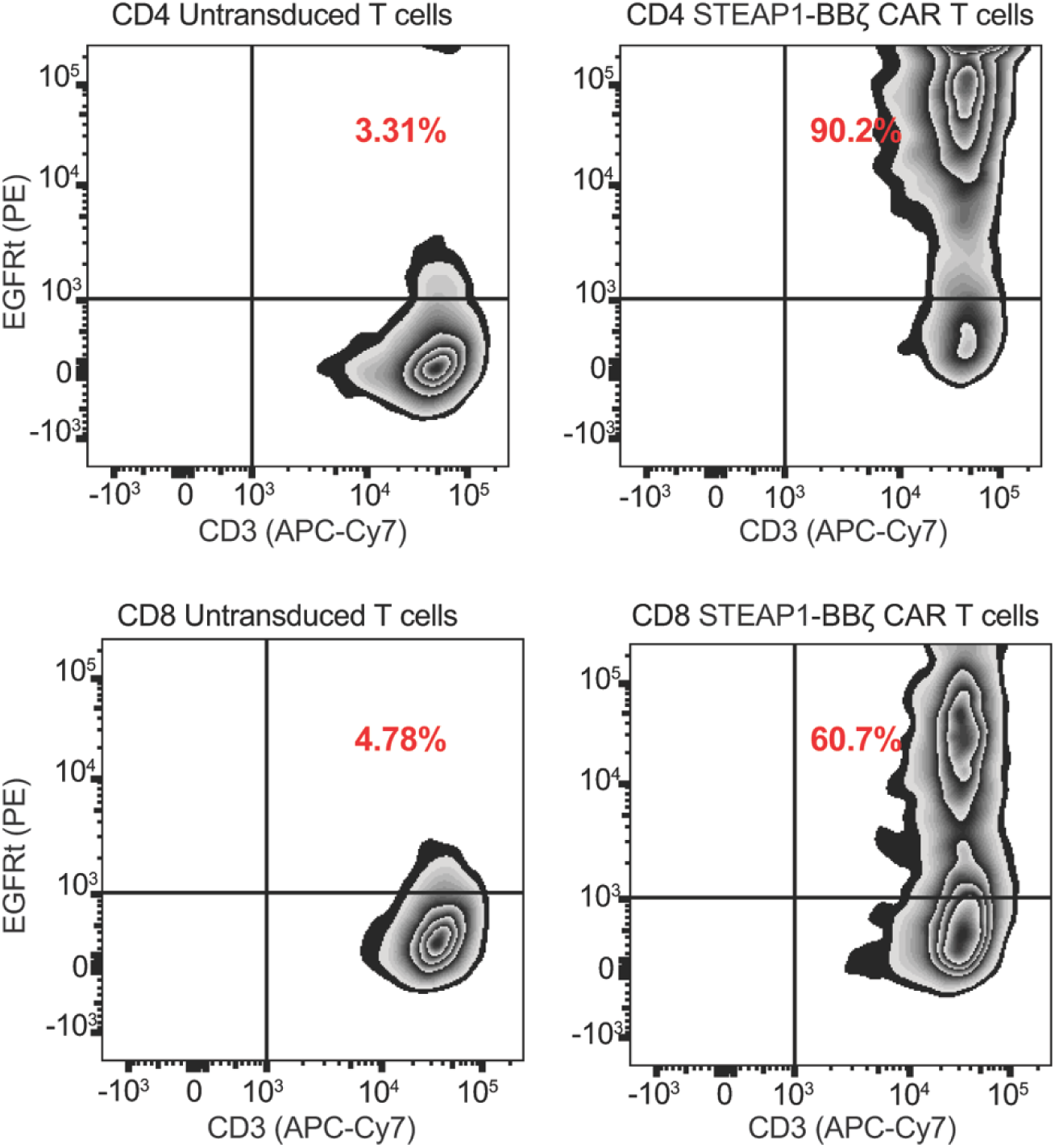
I**m**munophenotyping **of STEAP1-BB**ζ **CAR T cells.** Representative flow cytometry plots showing the immunophenotyping of CD3^+^ CD4^+^ and CD3^+^ CD8^+^ untransduced (EGFRt^-^) or STEAP1-BBζ (EGFRt^+^) T cells products at day 5 of expansion.

**Supplementary Figure 2.**
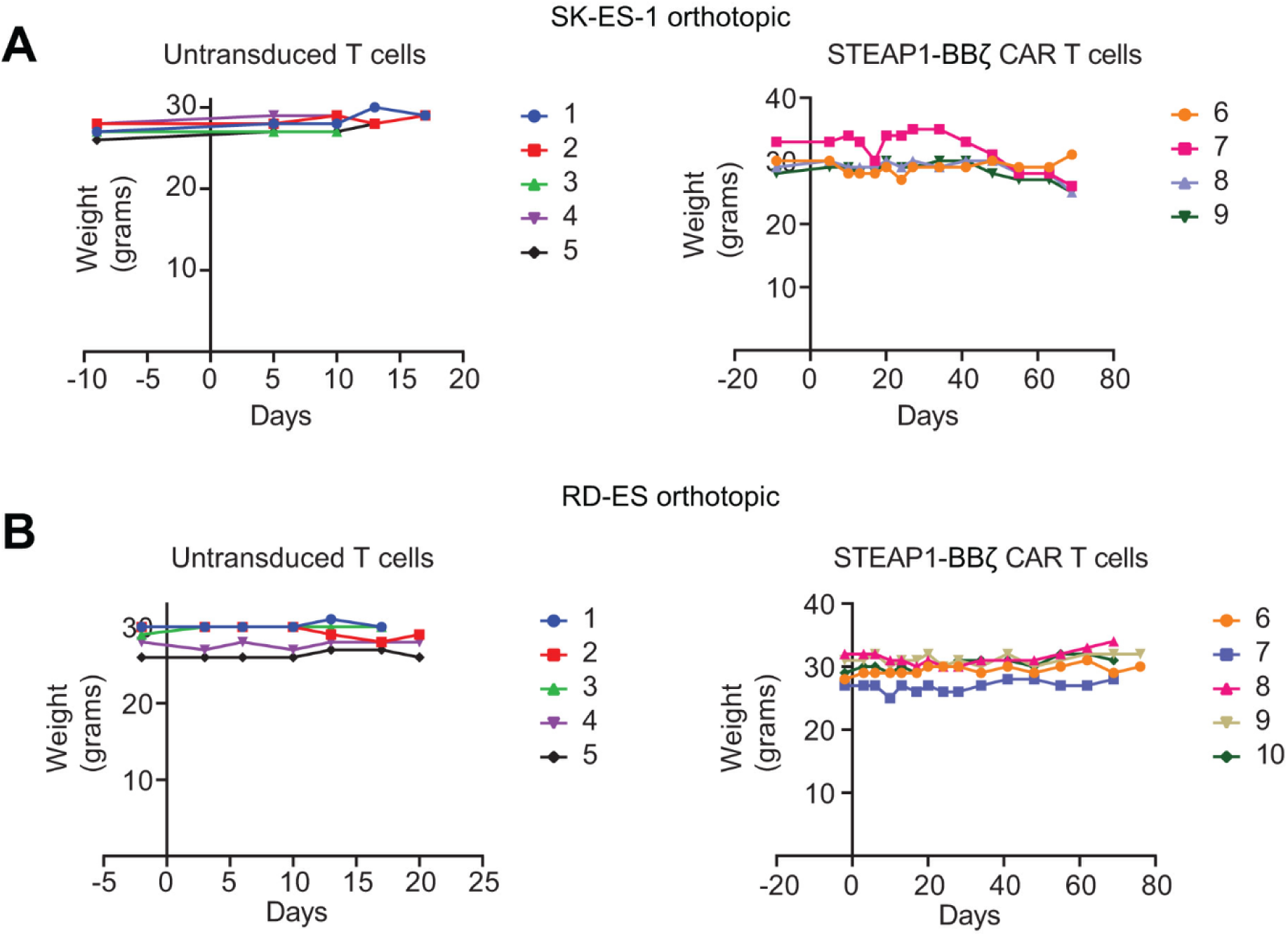
Body weight monitoring of mice treated with STEAP1-BB**ζ** CAR T cells in orthotopic EwS models. (A) Plots of individual body weight trajectories of mice bearing SK-ES-1 orthotopic tumors treated with either untransduced T cells or STEAP1-BBζ CAR T cells. (B) Plots of individual body weight trajectories of mice bearing RD-ES orthotopic tumors treated with either untransduced T cells or STEAP1-BBζ CAR T cells.

**Supplementary Figure 3.**
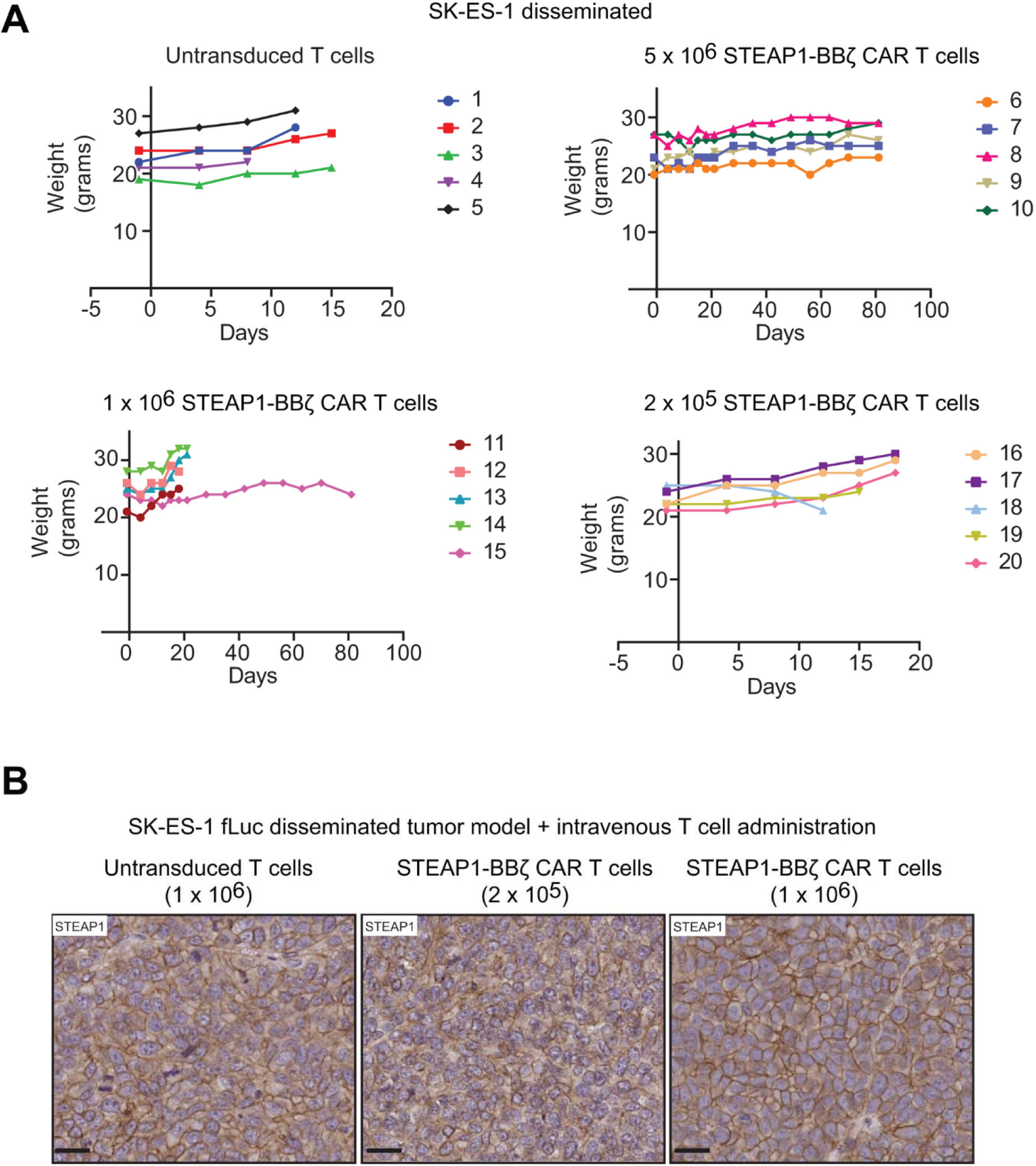
Body weight monitoring and STEAP1 antigen expression in mice treated with STEAP1-BB**ζ** CAR T cells in a disseminated EwS model. (A) Plots of individual body weight trajectories of mice bearing SK-ES-1 disseminated tumors treated with either untransduced T cells or different doses of STEAP1-BBζ CAR T cells. (B) Representative IHC images of STEAP1 expression in SK-ES-1 tumors following treatment with 2LxL10^5^ or 10^6^ STEAP1-BBζ CAR T cells showing retained STEAP1 expression post-treatment. Scale bars=50Lµm. STEAP1 IHC was performed on n=4 biologically independent tumors per group.

**Supplementary Figure 4.**
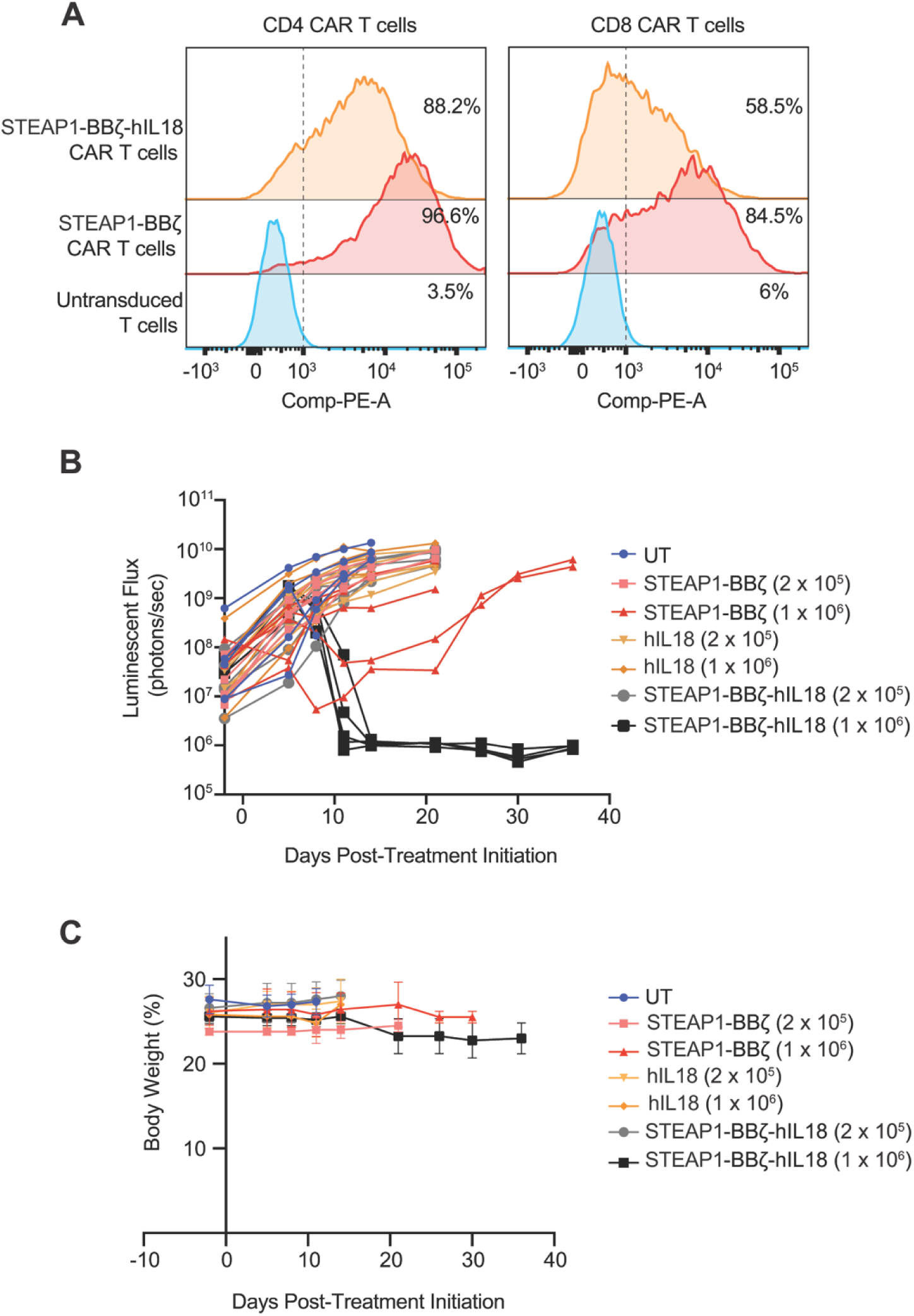
Immunophenotyping, tumor burden, and body weight analysis of STEAP1-BB**ζ**-hIL-18 CAR T cells in the SK-ES-1 disseminated EwS model. (A) Representative flow cytometry plots showing the immunophenotyping of CD4^+^ and CD8^+^ untransduced or STEAP1-BBζ or STEAP1-BBζ-hIL18 CAR T cells products at day 5 of expansion. (B) Quantification of total tumor flux over time from live BLI of mice treated with untransduced T cells, hIL-18T cells, STEAP1-BBζ CAR T cells, or STEAP1-BBζ-hIL-18 CAR T cells. Body weight trajectories of mice bearing SK-ES-1 disseminated tumors and treated with untransduced T cells, hIL-18T cells, STEAP1-BBζ CAR T cells, or STEAP1-BBζ-hIL-18 CAR T cells.

